# Complex interactions in legume/cereal intercropping system: role of root exudates in root-to-root communication

**DOI:** 10.1101/097584

**Authors:** Yinshan Jiao, Entao Wang, Wenfeng Chen, Donald L. Smith

**Affiliations:** State Key Laboratory of Agrobiotechnology, Beijing 100193, China; College of Biological Sciences and Rhizobia Research Center, China Agricultural University, Beijing 100193, China; Departamento de Microbiología, Escuela Nacional de Ciencias Biológicas, Instituto Politécnico Nacional, 11340 México D.F., Mexico; Department of Plant Science, Macdonald Campus of McGill University, Ste-Anne-de-Bellevue, Quebec, Canada. H9X 3V9

## Abstract

Dear Editor,

Legume/cereal intercropping systems have been regarded as the practical application of basic ecological principles such as diversity, competition and facilitation. In a recent PNAS paper, Li et al. (1) describe the novel finding that maize exudates promote faba bean nodulation and nitrogen fixation by upregulating genes involved in (iso)flavonoids synthesis (chalcone–flavanone isomerase) within faba bean, resulting in production of more genistein, a legume-to-rhizobia signal during establishment of the faba bean N_2_–fixing symbiosis. Although we salute the authors’ methodological efforts, there is another mechanism that could be responsible for the effect of corn root exudates on faba been nitrogen fixation observed in this article (1). The authors may misunderstand their data and the signalling role of maize exudates, thus got a defective model for the root interactions between faba bean and maize.

---

In their study, to explore the potential influence of maize exudates on the rhizobia physiological status, Li et al. (1) performed rhizobial growth curve by adding root exudates from maize and found no obvious affect. However, they did not check the possible effect of maize root exudates on the synthesis of Nod-factor. Previous data have showed that root washings and extracts from maize roots could directly induce the synthesis of Nod factor-like lipo-chitooligosaccharides (LCOs) of rhizobia in vitro (2). The amount of LCOs secreted by rhizobia cultured with root extracts from maize was even higher than those induced by soybean (host plant for the tested rhizobia) root extracts (2). In truth, the LCOs as key molecular recognized by legume host induce root hair deformation, infection thread formation and further trigger a series of symbiosis-related gene expression (3). It is likely that this mechanism also contributes to the observed increase in nodulation of faba bean. Therefore, future studies are needed to assess whether maize exudates may directly induce the rhizobia to produce more LCOs and enhance nodulation when interacted with faba bean.

Genistein, legume-specific isoflavonoids (4, 5), are signature characteristic of legumes (4, 5) and a key symbiotic signal in the *soybean-Bradyrhizobium* symbiosis (6, 7). However, Li et al. (1) found that the concentration of genistein in maize root exudates alone was similar to that of faba bean exudates alone (Fig. S4), which firstly evidenced that that genistein were synthesised by a nonlegume. This should be confirmed in future studies. Further, genistein was not detected in root exudates from a mixture of wheat and faba bean, but was present at high levels in exudates from faba bean alone (Fig. S4) (1), indicating possible suppression of gensitein production by faba bean roots by wheat root exudates.

Importantly, Li et al. (1) detected high expressions of some key genes of faba bean root after the addition of root exudates from maize compared to those of water-treated sample (Fig. 4). Among these genes, *NODL4* and *END93* were induced in 35-days faba bean root treatment with maize exudates, which was not consistent with the fact that early nodulin-like proteins have strong expression in early infection phase and nodule tissue (8). Additionally, *FixI* gene (GenBank no. KU973547) was described to encode a nitrogen fixation protein of faba bean and its expression could be detected in all root RNA samples (Fig. 4)(1). However, *FixI*, known as a member of bacterial fix cluster genes, involved in symbiotic nitrogen fixation in rhizobia (9) and should not be detected in plant root samples. To gain insight into the unidentified gene “*FixI*”, we blasted the submitted gene sequence in NCBI database. The result showed that this “*FixI*” gene has the highest similarity with an annotated heavy-metal-associated domain protein mRNA from *Medicago truncatula* (GenBank no. XM_003626494, 85% gene sequence identity), which has no genetic and molecular function information based on the available literature; it may be that this is a false “*FixI*” gene, and should not be used as an indicator of nitrogen fixation activity in faba bean roots.

To gain insight into if this false “*FixI*” gene in faba bean may have a function related to nitrogen fixation when symbiosis with rhizobia, we also analyzd the expression pattern of gene with most sequence similarity in *M. truncatula* (XM_003626494, mentioned above). It showed that the tested gene has highest expression in leaf compared with other organs and no clear inducted-expression in the root and nodule after inoculation with rhizobia in *M. truncatula* (Fig. 1). It means the false “*FixI*” gene could not involve in plant nitrogen fixation regulation.

**Fig. 1.**
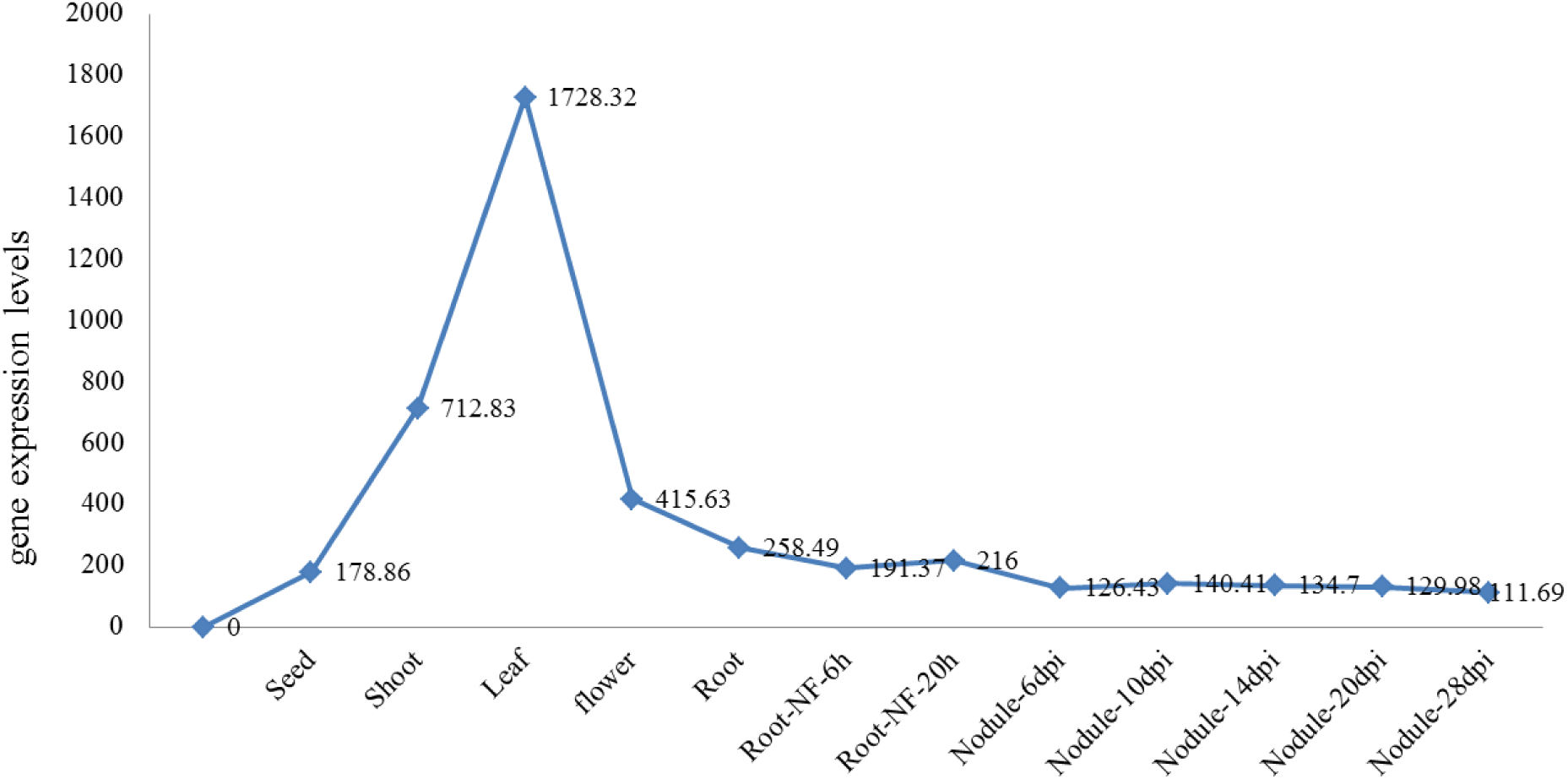
Expression analysis of an annotated heavy-metal-associated domain protein in *Medicago truncatula.* All used gene expression data were based on the Affymetrix GeneChip which server archives all publically-available *M. truncatula* gene expression data (http://bio_info.noble.org/gene-atlas/.). NF, nod factor.

It would seem that there are three potential mechanisms by which nodulation and N_2_ fixation can be increased by the root exudates in the legume-cereal intercropping systems: 1) reduced soil mineral N due to the cereal component, 2) enhanced interorganismal signalling due to the presence of appropriate (iso)flavonoids in the cereal root exudates and 3) induced production of (iso)flavonoid (genistein in the case of faba bean plants) following exposure to cereal (corn in this case) root exudates, as elucidated in the highly original findings of Li et al. (1).

Overall, the authors should probably collect additional molecular data to support their hypothesis and the potential contributing mechanisms indicated above should be noted. Although root exudates from maize may have essential factors in this facilitative effect, for example, we wait to see how these compounds help rhizobia to improve nodulation ability and enhance symbiosis of legume-rhizobium mutualism.

